# Suppression of MDA5-mediated antiviral immune responses by NSP8 of SARS-CoV-2

**DOI:** 10.1101/2020.08.12.247767

**Authors:** Ziwei Yang, Xiaolin Zhang, Fan Wang, Peihui Wang, Ersheng Kuang, Xiaojuan Li

**Affiliations:** Institute of Human Virology, Zhongshan School of Medicine, Sun Yat-Sen University, Guangzhou, Guangdong, 510080, China; Key Laboratory of Tropical Disease Control (Sun Yat-Sen University), Ministry of Education, Guangzhou, Guangdong, 510080, China; Advanced Medical Research Institute, Shandong University, Jinan, Shandong 250012, China

**Keywords:** SARS-CoV-2, NSP8, MDA5, interferon, immune evasion

## Abstract

Melanoma differentiation-associated gene-5 (MDA5) acts as a cytoplasmic RNA sensor to detect viral dsRNA and mediates type I interferon (IFN) signaling and antiviral innate immune responses to infection by RNA viruses. Upon recognition of viral dsRNA, MDA5 is activated with K63-linked polyubiquitination and then triggers the recruitment of MAVS and activation of TBK1 and IKK, subsequently leading to IRF3 and NF-κB phosphorylation. Great numbers of symptomatic and severe infections of SARS-CoV-2 are spreading worldwide, and the poor efficacy of treatment with type I interferon and antiviral agents indicates that SARS-CoV-2 escapes from antiviral immune responses via an unknown mechanism. Here, we report that SARS-CoV-2 nonstructural protein 8 (NSP8) acts as an innate immune suppressor and inhibits type I IFN signaling to promote infection of RNA viruses. It downregulates the expression of type I IFNs, IFN-stimulated genes and proinflammatory cytokines by binding to MDA5 and impairing its K63-linked polyubiquitination. Our findings reveal that NSP8 mediates innate immune evasion during SARS-CoV-2 infection and may serve as a potential target for future therapeutics for SARS-CoV-2 infectious diseases.

**Importance:** The large-scale spread of COVID-19 is causing mass casualties worldwide, and the failure of antiviral immune treatment suggests immune evasion. It has been reported that several nonstructural proteins of severe coronaviruses suppress antiviral immune responses; however, the immune suppression mechanism of SARS-CoV-2 remains unknown. Here, we revealed that NSP8 protein of SARS-CoV-2 directly blocks the activation of the cytosolic viral dsRNA sensor MDA5 and significantly downregulates antiviral immune responses. Our study contributes to our understanding of the direct immune evasion mechanism of SARS-CoV-2 by showing that NSP8 suppresses the most upstream sensor of innate immune responses involved in the recognition of viral dsRNA.

## Introduction

Severe acute respiratory syndrome coronavirus 2 (SARS-CoV-2) is an emerging severe coronavirus that is currently causing a global outbreak of Coronavirus Disease 2019 (COVID-19). It has infected more than twenty million patients and caused more than two-thirds of a million deaths. The number of patients and deaths are still rapidly increasing; however, no effective therapy, vaccine or cure are available. Knowledge of SARS-CoV-2 infection, pathogenesis, diseases and treatments remains limited, and novel targets of therapeutics and drug development are quite urgently needed.

Analysis of clinical data from different SARS-CoV-2 patients has shown that excessive cytokine release, known as a “cytokine storm”, is closely related to disease severity ^1–3^. To characterize the host immune and inflammatory responses in COVID-19 patients, genome-wide RNA-sequencing analysis was performed, which indicated that the proportion of immune cells in the blood was reduced in patients who required non-ICU admission, with lower levels of *GCSF, CXCL10*/*IP-10, CCL2*/*MCP-1* and *CCL3*/*MIP-1A* detected. In addition, some anti-inflammatory cytokines, such as IL-10 and TGF-β, were found to be induced during SARS-CoV-2 infection ^4^. Accordingly, compared with ICU care patients, non-ICU care patients had lower plasma levels of cytokines, including IL-2, IL-7, IL-10, GSCF, IP10, MCP1, MIP1A, and TNFα. Furthermore, mild cases of COVID-19 exhibited decreased plasma levels of IL2R and IL6 compared with severe cases, while an excessive inflammatory response was observed in dead cases ^5,6^. These studies reported a correlation between the release of inflammatory cytokines and pathogenesis of SARS-CoV-2.

SARS-CoV-2 and other coronaviruses generate massive amounts of RNA products during their infection that are then recognized by host cytosolic RNA sensors, including retinoic acid-inducible gene I (RIG-I) and melanoma differentiation-associated gene-5 (MDA5) ^7^. Activation of RIG-I and MDA5 triggers the formation of signalosomes that induce the expression of type I IFN and ISGs and the subsequent execution of an antiviral state within the cell ^8^. To escape immune elimination and to survive and then replicate, coronaviruses, including SARS-CoV and MERS-CoV, have evolved strategies to inhibit or delay IFN production and responses. SARS-CoV encodes a set of accessory proteins, several of which target the innate immune response. Its nonstructural protein 1 (NSP1) binds to the 40S ribosome to inactivate translation and induces host mRNA degradation ^9,10^; NSP15 was identified as an IFN antagonist to enhance immune evasion ^11,12^; NSP16, a 2′O-methyltransferase (2′O-MTase), provides a cap structure at the 5′-end of viral mRNAs to evade MDA5 detection ^13–15^; furthermore, open reading frame 6 (ORF6) also reduces IFN production ^16^. In addition, ORF-9b localizes to mitochondrial membranes to induce the degradation of MAVS, TRAF3 and TRAF6, severely limiting host cell IFN production ^17^. In addition, MERS ORF4b is involved in evasion of the innate immune response by binding α-karyopherin proteins, leading to the inhibition of NF-κB nuclear translocation ^18^.

Considering the numerous existing asymptomatic infected populations, it is reasonable to suggest that SARS-CoV-2 has evolved effective strategies to inhibit the host immune response, which is also a challenge for preventing and treating COVID-19. Consistent with these observations, a recent study showed that NSP1 of SARS-CoV-2 shuts down host mRNA translation by binding to 40S and 80S ribosomes, effectively blocking RIG-I-dependent innate immune responses ^19^. However, the potential immune inhibitory mechanism mediated by viral components of SARS-CoV-2 is not well known.

We performed a systematic screening and determined that SARS-CoV-2 NSP8 is a suppressor of the type I immune response upon RNA viral infection. It decreases type I IFN production and ISG expression, by which it interacts with the MDA5 CARD domain and then inhibits MDA5 K63-linked polyubiquitination and thus terminates MDA5-mediated immune responses.

## Results

### NSP8 inhibits viral RNA-related type I interferon and antiviral responses

To investigate a factor of SARS-CoV-2 that suppresses the type I IFN signaling pathway, we screened a panel of viral nonstructural proteins (NSPs) that regulate the expression of interferon and inflammatory factors using RT-PCR arrays. 293T cells were transfected with empty vector or NSP-expressing plasmids for 24 h. The cells were left untreated or challenged with dsRNA analog poly(I:C) for 18 h or 24 h separately and then harvested for analysis of gene expression. Several NSPs exhibited the inhibitory effects on immune and inflammatory gene expression (Supplementary Fig.1). We found that NSP8 significantly downregulated the expression of *TNF-α, IFN-β*, interferon-stimulated genes (ISGs) *IFIT1* and *IFIT2* and proinflammatory cytokines *IL-6* and *CCL-20* (Fig.1a). Consistent with our results, NSP1 was previously reported to inhibit the translation of type I interferon to evade the immune response, which validated our screening approach ^9,19^.

**Figure 1.**
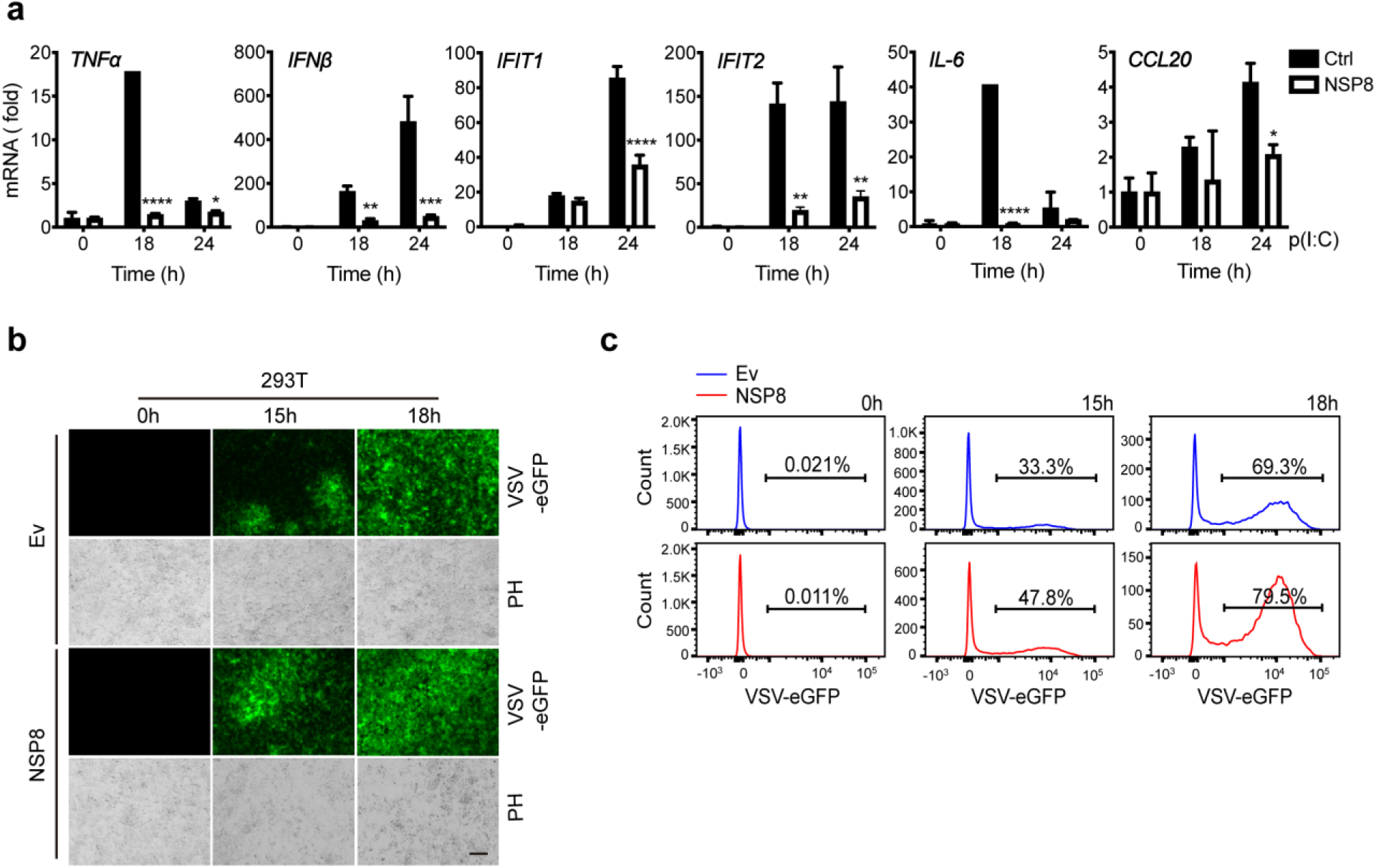
NSP8 inhibits the viral RNA-induced type I IFN signaling pathway. a. HEK293T cells were transfected with a control vector (Ctrl) or *Flag-NSP* plasmids. Twenty-four hours post transfection, cells were treated with poly(I:C) (5 μM) for the indicated time points and then subjected to RT-PCR analysis for *TNF-α, IFN-β, IFIT1, IFIT2, IL-6* and *CCL20* expression. Data are shown as the mean values ± SD (n = 3). *, *p* < 0.0332; **, p < 0.0021; ***, p < 0.0002; ****, p < 0.0001; by Sidak’s multiple comparisons test. b. HEK293T cells were transfected with an empty vector or *Flag-NSP8*. Twenty-four hours post transfection, cells were infected with VSV-eGFP (MOI = 0.01) for the indicated time points and subjected to phase-contrast (PH) or fluorescence analyses. Scale bar, 15 μm. c. eGFP-positive cells were detected by flow cytometry. Numbers indicate the representative percentage of eGFP-positive cells of three independent experiments.

To confirm that the attenuation of the type I IFN response by NSP8 is correlated with a decrease in the antiviral response, *NSP8-*overexpressing 293T cells were subsequently infected with vesicular stomatitis virus tagged with enhanced GFP (VSV-eGFP). As analyzed by fluorescence microscopy and flow cytometry, NSP8 overexpression increased the percentage of GFP-positive cells with the extended response time compared with empty vector-transfected cells (Fig.1b,c), suggesting that NSP8 promoted the efficiency of viral infection and gene expression, hence favoring VSV-eGFP replication. These data demonstrated that NSP8 is an inhibitory protein in type I IFN signaling and antiviral responses during SARS-CoV-2 infection.

### NSP8 decreases MAVS-dependent antiviral responses

To further understand whether the inhibitory effects of NSP8 are exerted on both the IRF3 and NF-κB pathways, 293T cells were cotransfected with empty vector or IRF3 or p65 expressing plasmid and with NSP8-expressing plasmid, followed by VSV infection for the indicated time points. By immunoblotting analysis to detect the effect of NSP8 on the phosphorylation of TBK1 (pTBK1), the phosphorylation of IRF3 (pIRF3) (Fig.2a), the phosphorylation of IKKα/β (pIKKα/β), the phosphorylation of p65 (p-p65) and IκBα (Fig.2b), we found that in NSP8-overexpressing cells, pTBK1 level was four- to nine-fold lower than that in control cells after VSV infection. As a downstream transcription factor that is phosphorylated and activated by TBK1, IRF3 phosphorylation was entirely blocked in NSP8-overexpressing cells (Fig.2a). Similarly, inhibition of IKKα/β phosphorylation by NSP8 led to the stabilization of IκBα, and NF-κB signaling was strongly inhibited by NSP8 overexpression, as indicated by the decrease in p65 phosphorylation (Fig.2b). These results confirm that NSP8 suppresses the activation of both IRF3 and NFκB and the upstream cascade of the type I IFN signaling pathway.

**Figure 2.**
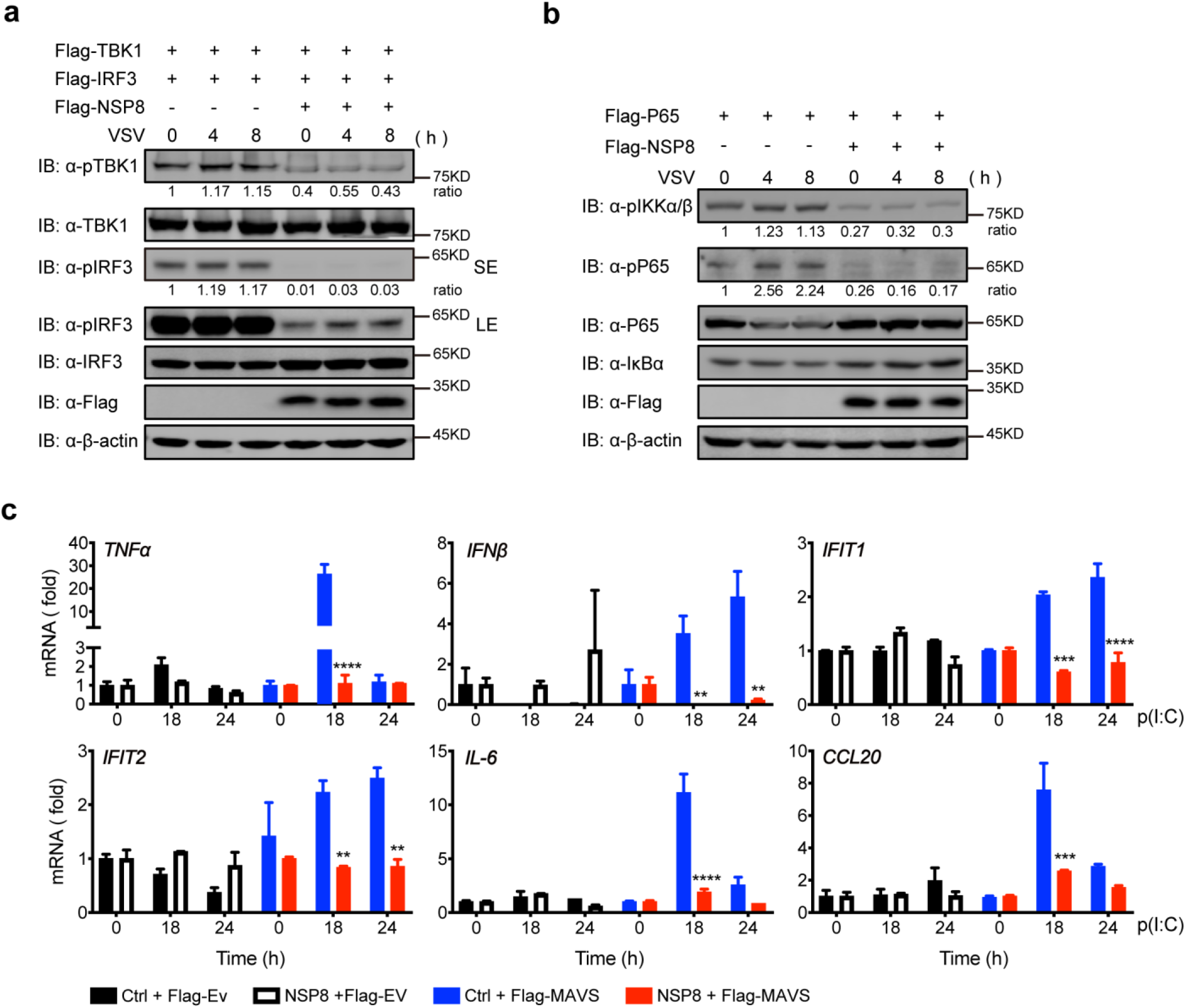
NSP8 inhibits the MAVS-dependent immune signaling pathway. a-b. HEK293T cells were transfected with an empty vector or *Flag-NSP8* for 8 h and then transfected with *Flag-TBK1* and *Flag-IRF3* (a) or transfected with *Flag-p65* (b). Twenty-four hours post transfection, cells were infected with VSV-eGFP (MOI = 1) for the indicated time points and then harvested. Whole cell extracts were analyzed by immunoblotting as indicated. Representative images of three independent experiments are shown. c. *MAVS* knockout HEK293T cells were cotransfected with *Flag-NSP8* and an empty vector or *Flag-MAVS*. Twenty-four hours post transfection, cells were treated with poly(I:C) (5 μg/ml) for the indicated time points, and then total RNA was extracted and subjected to RT-PCR analysis for *TNF-α, IFN-β, IFIT1, IFIT2, IL-6* and *CCL20* expression. The data are shown as the mean values ± SD (n = 3). *, p < 0.0332; **, p < 0.0021; ***, p < 0.0002; ****, p < 0.0001; by Sidak’s multiple comparisons test.

To investigate whether NSP8 plays a role in the early step of the type I IFN signaling cascade, we detected the inhibitory effect of NSP8 in MAVS-deficient vs. complemented cells. *MAVS* knockout HEK293T cells (HEK293T^MAVS-/-^) were cotransfected with *NSP8*-expressing plasmid or empty vector and with *MAVS-* expressing plasmid or empty vector and then stimulated with poly(I:C) for different times. As expected, the expression of *TNF-α, IFN-β, IFIT1, IFIT2, IL-6* and *CCL-20* was not downregulated by NSP8 upon stimulation (Fig.2c, black vs. white columns), while MAVS overexpression in HEK293T^MAVS-/-^ cells significantly restored their induction upon stimulation, and NSP8 once again had suppressive effects on the expression of these cytokines (Fig.2c, red vs. blue columns). Taken together, these results suggest that NSP8 suppresses MAVS-dependent innate immune responses, probably by acting on either MAVS or upstream RNA sensors.

### NSP8 interacts with MDA5 and suppresses MDA5 K63-linked polyubiquitination

Since viral RNA of coronaviruses contains a methylated 5’-cap and 3’-polyA tail that is similar to cellular mRNA, we assumed that NSP8 may preferentially regulate MDA5-mediated responses rather than RIG-I-mediated responses that recognize 5’-pppRNA. Hence, we performed coimmunoprecipitation (co-IP) analysis and found that NSP8 interacts with MDA5 (Fig.3a). Furthermore, we mapped the binding domain and determined that its CARD domain is responsible for the NSP8 interaction (Fig.3b). As a result, MDA5-mediated ISRE-luc activity was inhibited by NSP8 in a dose-dependent manner (Fig.3c). In addition, confocal microscopy demonstrated that NSP8 tightly colocalized with MDA5 inside cells (Fig.3d). These results suggest that NSP8 interacts with MDA5 and directly suppresses MDA5-mediated immune responses.

**Figure 3.**
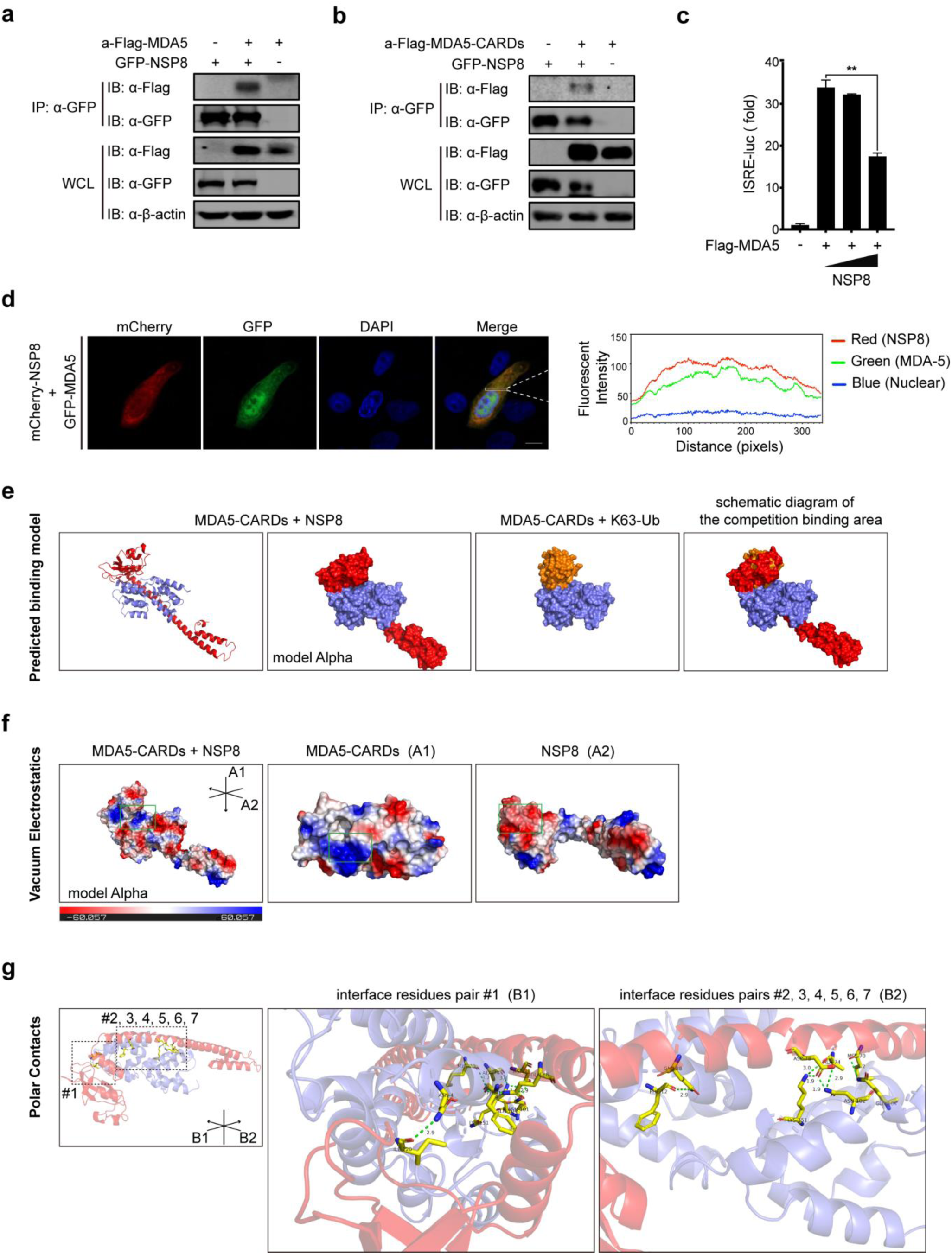
NSP8 interacts with MDA5 at the CARD domain. a-b. HEK293T cells were transfected with *GFP-NSP8* and full-length *Flag-MDA5* (a) or *Flag-MDA5-CARD* (b), and after 36 h, the cells were collected and lysed. Then, cell lysates were subjected to coimmunoprecipitation using anti-GFP beads, followed by immunoblotting with the indicated antibodies. c. HEK293T cells were transfected with an empty vector or increasing amounts of *NSP8-*expressing plasmid plus an *ISRE-luc* reporter and *MDA5*-expressing plasmids. Twenty-four hours post transfection, the cells were collected, and then cell lysates were analyzed for ISRE-luc activit*y.* The representative results of three independent experiments are shown as the mean values ± SD (n = 3). d. HEK293T cells were cotransfected with mCherry-NSP8 and GFP-MDA5. Twenty-four hours post transfection, the images were visualized by confocal microcopy analysis. Scale bar, 10 μm. The colocalization coefficient was determined by qualitative analysis of the fluorescence intensity of the selected area in Merge. e-g. Computer-based prediction and structural modeling of the interaction of NSP8-MDA5 CARD domains. e. Structural prediction of the MDA5-CARD-NSP8 interaction and MDA5-CARD linkage with K63-Ub. PDB structures were input into ZDOCK Server for docking calculation separately. The predicted binding models of MDA5 CARDs with NSP8 and MDA5 CARDs with K63-Ub were processed in PyMOL for demonstration. Model alpha simulates the protein surface. Red chain, NSP8; violet chain, MDA5 CARDs; brown chain, K63-Ub. f. Model alpha in (e) was subjected to vacuum electrostatics calculation in PyMOL. A1 and A2 indicate the viewing angle in the green frame. The green frame indicates the contact area demonstrated in A1 and A2. Scale bar indicates the range of vacuum electrostatics. g. Polar contacts within interface of MDA5-CARD-USP8 were demonstrated with PyMOL. B1, B2 indicates the viewing angle of dashed borders. #1, 2, 3, 4, 5, 6, 7 indicates paired residues. Paired residues were highlighted with sticks model in yellow color. Green dashed lines indicate polar contacts between paired residues, number besides dashed line indicates distance between two atoms connected (Å).

To further understand the molecular mechanism by which NSP8 interacts and interferes with MDA5, we predicted the MDA5 CARD, NSP8 and K63-Ub tertiary structures with SWISS-MODEL, and then the predicted structures were input into ZDOCK SERVER for simulation. Predicted docking models were processed in PyMOL for visualization. Surprisingly, we found that NSP8 possesses a long α-helix (Supplementary Fig.2a), which is tightly packed in the ravines formed by the two α-helixes of MDA5 CARDs. The random coil and a short α-helix in the N terminus of NSP8 occupy the area or space that interacts with K63-Ub (Fig.3e and Supplementary Fig.2b). Further calculation of vacuum electrostatics for this binding model demonstrated that the contact area in the chain of MDA5 CARDs is positively charged, while the corresponding area in the chain of NSP8 is negatively charged (Fig.3f and Supplementary Fig.2c), implying that there is a likely interaction of these two structures. We further searched the polar contacts in the interface of the binding model with PyMOL and found that 7 paired residues anchor with each other, one locates in the N-terminal coil of NSP8 while the others are in the long α-helix (Fig.3g).Thus, computer-based molecular structural prediction and modeling implies that NSP8 interacts with the MDA5 CARD domain probably through ionic interactions and dipolar surfaces between the NSP8 and MDA5 CARD binding pockets.

Next, we sought to determine how NSP8 inhibits MDA5 activation. It is well documented that upon virus infection, the MDA5 CARD domain undergoes K63-linked polyubiquitination and recruits MAVS to form a signalosome ^20^. The structural prediction of the NSP8-MDA5 CARD interaction showed that NSP8 may interrupt this process since it interacts with MDA5 at its CARD domain and shields the binding area or space for K63-ubiquitin linkage (Fig.3e, Supplementary Fig.2, and pdb file). The polyubiquitination of MDA5 was analyzed in the presence or absence of NSP8 expression. The *MDA5*-expressing plasmid was cotransfected into HEK293T cells with a WT-, K48-, or K63-linked ubiquitin-expressing plasmid, and an in vivo ubiquitination assay showed that MDA5 WT- and K63-linked polyubiquitination were strongly inhibited (Fig.4a,c), while K48-linked polyubiquitination was barely affected (Fig.4b). Thus, these results reveal that NSP8 interferes with the MDA5-MAVS signalosome by inhibiting the K63-linked polyubiquitination of MDA5.

**Figure 4.**
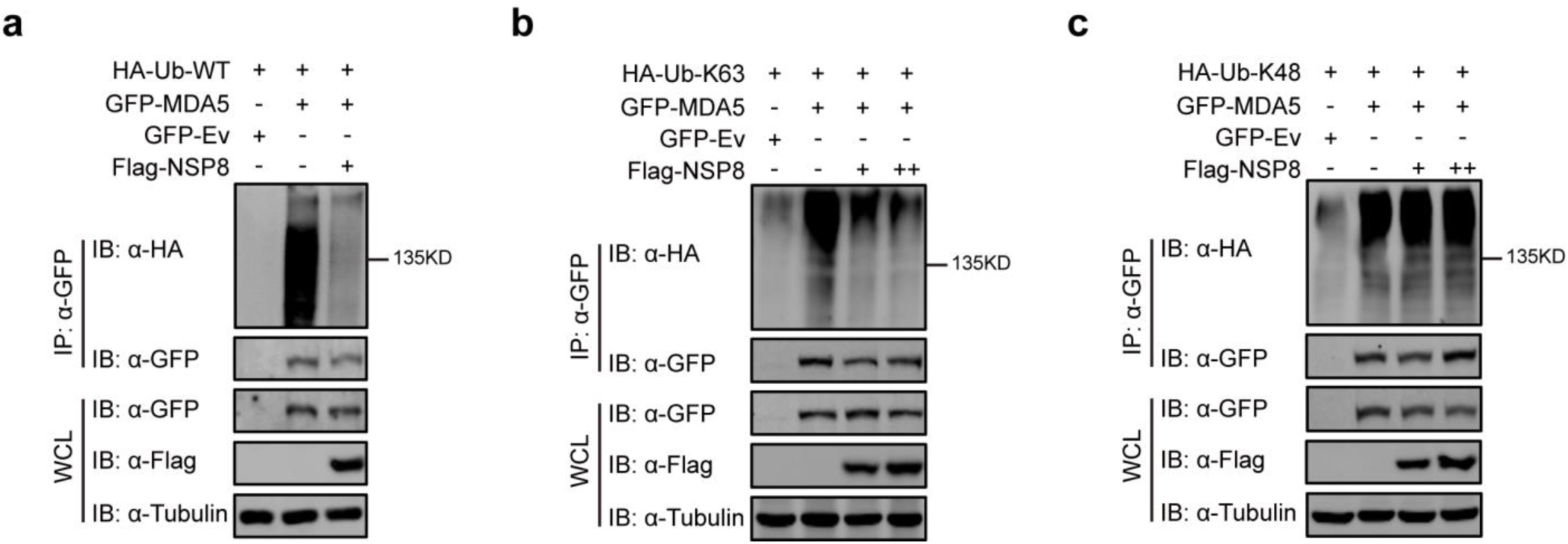
NSP8 suppresses K63-linked polyubiquitination of MDA5. a-c. HEK293T cells were cotransfected with *HA-Ub-WT* (a), *HA-Ub-K63* (b) or *HA-Ub-K48* (c) and *GFP-MDA5* or *GFP-*tagged empty vector plus *Flag-NSP8.* Twenty hours post transfection, cells were treated with MG132 (5 μM) for 4 h, and then cells were collected and lysed in 0.1% SDS-containing lysis buffer. Cell lysates were subjected to coimmunoprecipitation using anti-GFP beads, followed by immunoblotting analysis with the indicated antibodies. Representative images of three independent experiments are shown.

### NSP8 decreases the expression of antiviral immune and inflammatory genes

Next, we investigated whether NSP8 affects the downstream expression of immune and inflammatory genes and cytokines. NSP8-expressing A549 cells were collected, and total RNA was subjected to RT-PCR array analysis for immune and inflammatory gene expression (Primer listed in Supplementary Table.1). The expression of the majority of cytokines, including *IL-1β, IL-2, IL-5, IL-6, IL-26, IL-33, IFN-β, IFIT1* and *IFIT2*, was downregulated by NSP8 expression. The transcription of some pleiotropic chemoattractant cytokines, such as *IL-16, IL-17A, IL-17F*, and *IL17C*, was also downregulated. Furthermore, a decreasing transcription tendency was also observed for the inflammatory receptors *IL-1RI, IL-1RII, IL-2Rα*, and *IL18RII*; NK cell-associated activation receptors, such as *NKp44, NKp46*, and *NKG2B*; and the trans-acting T-cell-specific transcription factor *GATA3* (Fig.5a), indicating that the activation of T cells and NK cells was attenuated by NSP8 through the suppression of these key factors. Although these decreased cytokines and receptors may not be directly activated by IRF3 or NFκB, they could be regulated by downstream cytokines or other factors derived from these two pathways. In contrast, the cytokine IL-2 and IFN-gamma suppression gene *FOXP3* was significantly increased with NSP8 overexpression.

**Figure 5.**
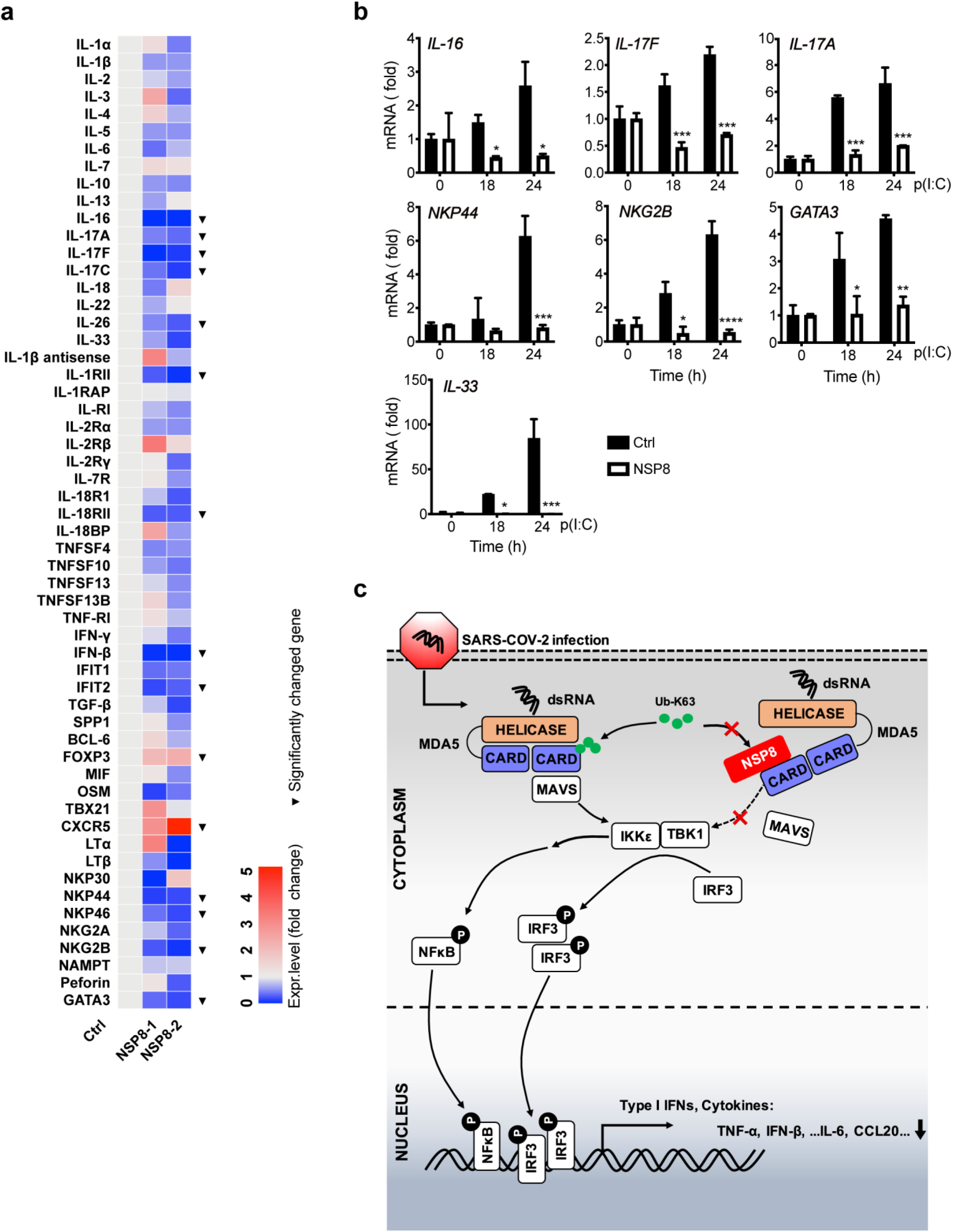
NSP8 decreases the expression of antiviral immune and inflammatory factors. a. A549 cells were transfected with an empty vector or *Flag-NSP8*. Thirty-six hours post transfection, cells were collected, and total RNA was extracted and subjected to RT-PCR analysis for the expression of the indicated genes. “▾” indicates genes that were significantly changed in two independent *NSP8-*expressing samples. b. A549 cells were transfected with an empty vector or *Flag-NSP8*. Twenty-four hours post transfection, cells were treated with poly(I:C) (5 μg/ml) for the indicated time points, and then total RNA was extracted and subjected to RT-PCR analysis of the expression of selected genes. The data are shown as the mean values ± SD (n = 3). *, p < 0.0332; **, p < 0.0021; ***, p < 0.0002; ****, p < 0.0001; by Sidak’s multiple comparisons test. c. Working model of SARS-CoV-2 NSP8 negatively regulating the MDA5-mediated type I IFN signaling pathway. NSP8 interacts with MDA5 and impairs its K63-linked polyubiquitination, thus inhibiting the phosphorylation of TBK1 and IRF3 and subsequently downregulating the production of type I IFNs, immune cytokines and inflammatory factors.

To further confirm that the downregulation of these immune and inflammatory cytokines and genes is mediated by NSP8 under physiological conditions, A549 cells were transfected with an NSP8-expressing plasmid or empty vector and then stimulated with poly(I:C) mimicking viral RNA for the indicated times. The inhibition of the expression of key cytokines and related genes was verified, and NSP8 negatively regulated the expression of these immune and inflammatory genes (Fig.5b). Collectively, these results suggest that NSP8 could strongly impair the expression of genes involved in antiviral immune and inflammatory responses.

## Discussion

It has not been well investigated whether SARS-CoV-2 components have immunosuppressive functions, so we are very interested in exploring the roles of SARS-CoV-2 proteins, especially the nonstructural protein family (NSP), in viral immune evasion. In the present study, we screened a panel of NSPs and identified NSP8 as an immune suppressor of the type I IFN signaling pathway. We have shown that NSP8 overexpression impaired type I IFN production and the gene expression of immune and inflammatory factors, including *TNF-α, IFN-β, IFIT1, IFIT2, IL-6* and *CCL-20*. As a consequence, we observed that VSV-eGFP infection was significantly increased in cells with NSP8 overexpression. The subsequent detection of phosphorylation of kinases TBK1 and IKKα/β and substrates IRF3 and p65 provided evidence that both IRF3 and NFκB activity were inhibited by NSP8. MAVS is an upstream regulator of the type I IFN signaling cascade and regulates both pathways; hence, we overexpressed NSP8 in MAVS knockout HEK293T cells and observed that the inhibitory function of NSP8 towards type I IFN and cytokines was completely abolished. Re-expressing MAVS in MAVS KO HEK293T cells that also express NSP8 successfully restored the NSP8 inhibitory activity. Consequently, a series of antiviral immune and inflammatory cytokines and related genes were further strongly downregulated by NSP8 expression.

Herein, we speculated that NSP8 may act on RIG-I or MDA5, two upstream viral RNA sensors. Our results showed that NSP8 directly interacted with MDA5 on its CARD domains, and MDA5-mediated type I IFN signaling activities were strongly inhibited at the same time. Polyubiquitinated modification of MDA5 is crucial for its antiviral responses, K48-linked polyubiquitination mediates MDA5 proteasomal degradation ^21^, and K63-linked polyubiquitination mediates MDA5-induced type I IFN expression ^20^. We speculated that NSP8 may inhibit type I IFN signaling through the polyubiquitinated modification of MDA5. To test our hypothesis, we determined the status of WT-, K48- and K63-linked polyubiquitination of MDA5 in the absence or presence of NSP8 and confirmed that WT- and K63-linked polyubiquitination were impaired by NSP8, while K48-linked polyubiquitination was barely changed. We therefore conclude that under certain circumstances, NSP8 jeopardizes antiviral responses by impairing MDA5 K63-linked polyubiquitination.

Based on our existing experimental data, we propose a simple working model to illustrate how NSP8 negatively regulates innate immune responses by inhibiting MDA5 K63-linked polyubiquitination (Fig.5c). Upon SARS-CoV-2 infection, cytosolic viral dsRNA is recognized by MDA5 and triggers type I IFN expression; meanwhile, NSP8 is largely expressed and localized in the cytoplasm of host cells and then interacts with MDA5 on its CARD domains to inhibit its K63-linked polyubiquitination, consequently negatively regulating type I IFN signaling and antiviral immune responses, thus favoring viral infection and replication.

Interestingly, a recent study showed that in the carrier and intermediate host of coronaviruses—pangolin— the MDA5 gene is mutated and dysfunctional ^22^, while RIG-I and TLR 3, 7 and 8 are conserved among three different species. Considering the coexisting relationship between pangolins and coronaviruses and the tolerance of pangolins for SARS-CoV-2 infection, this finding implied that coronaviruses may be inclined to silence MDA5-mediated innate immunity because they could generate more drastic reactivity and injury upon viral RNA recognition. Based on this observation and the findings of our present study, the suppressive function of NSP8 in antiviral responses and its impairment of MDA5 activation have been identified, representing a direct immune evasion mechanism of the recognition of viral dsRNA.

Multiple posttranslational modifications, including phosphorylation, acetylation, methylation and polyubiquitination, are employed to regulate the antiviral signalosome ^23^. Among these modifications, polyubiquitination is commonly used for the degradation or activation of MDA5 and RIG-I. The E3 ubiquitin ligase TRIM40 targets MDA5 and RIG-I to promote their K27- and K48-linked ubiquitination, thus leading to their proteasomal degradation for immune silencing, but upon RNA virus infection, TRIM40 is downregulated to allow the activation of a sufficient antiviral immune response ^21^. Likewise, some viral proteins, such as Epstein-Barr virus protein BPLF1, which has ubiquitin- and NEDD8-specific deconjugase activity, interact with scaffold proteins 14-3-3 and TRIM25 to form a tri-molecular complex, consequently promoting the dimerization and ubiquitination of TRIM25. Consequently, K63-linked polyubiquitination of RIG-I is downregulated, leading to the attenuation of RIG-I-mediated type I IFN antiviral responses ^24^. In addition, the NS3 protein of ZIKA virus interacts with scaffold proteins 14-3-3ϵ and η separately through its 14-3-3 binding motif, hence blocking the translocation of RIG-I and MDA5 from the cytosol to mitochondria, impairing signalosome formation with MAVS, and antagonizing innate immunity ^25^. Our studies revealed that NSP8 of SARS-CoV-2 acts as a binding partner of MDA5 to shield its K63-linked polyubiquitination and then impairs the formation or activation of the MDA5 signalosome.

In summary, our study provides insights into the potential mechanisms of SARS-CoV-2 NSP8 in the inhibition of type I IFN signaling and antiviral responses. We provide compelling evidence that NSP8 plays a critical negative role in MDA5-mediated antiviral responses and demonstrate specific orchestration of the viral dsRNA-triggered signalosome and signal cascade by NSP8. Importantly, considering that MDA5 plays a key pathological role in antiviral immunity towards severe coronaviruses, antagonists of NSP8 could serve as a promising therapeutic target for COVID-19 therapies.

## Materials and Methods

### Cell culture and antibodies

HEK293T cells were cultured in DMEM (Gibco) supplemented with 10% fetal bovine serum (FBS). A549 cells (human adenocarcinoma lung tissue-derived epithelial cells) were cultured in RPMI 1640 (Gibco) medium containing 10% FBS. The following antibodies were used in this study: goat anti-mouse IRDye680RD (C90710-09) and goat anti-rabbit IRDye800CW (C80925-05), which were purchased from Li-COR; anti-HA (M132-3), purchased from MBL; anti-Flag (AE005), anti-GFP (AE0122) and anti-IRF3 (A2172), purchased from ABclonal; anti-β-actin (HC201-01), purchased from TransGen; and anti-pTBK1 (#5483P), anti-TBK1 (#3504), anti-pIRF3 (#4947), anti-phospho-NF-κB p65 (#3033), and anti-NF-κB p65 (#8242), purchased from Cell Signaling Technology.

### Plasmids

The following plasmids were used. Flag-NSP8 was kindly provided by the Peihui Wang lab (Shandong University). HA-tagged Ub, K48-Ub (K48 only), and K63-Ub (K63 only) were kindly provided by Dr Yang Du (Sun Yat-Sen University). To generate GFP- and mCherry-tagged NSP8 and GFP- and Flag-tagged MDA5, the NSP8 and MDA5 fragments were subcloned into the pEGFP-C2, p-mCherry-C2 and pcDNA3.1 vectors.

### Transfection and luciferase reporter assays

HEK293T cells were seeded in 24-well plates overnight and then transfected using Lipofectamine 2000 (Invitrogen) with 100 ng ISRE luciferase reporter (firefly luciferase), 20 ng pRL-TK plasmid (Renilla luciferase), 150 ng Flag-MDA5 expressing plasmid and increasing amounts (0, 100, or 200 ng) of NSP8-expressing plasmid. Twenty-four hours post transfection, cells were collected, and luciferase activity was measured with a Dual-Luciferase Assay kit (Promega) with a Synergy2 Reader (Bio-Tek) according to the manufacturer’s protocol. The relative level of gene expression was determined by normalization of firefly luciferase activity to Renilla luciferase activity.

### Virus infection

VSV-eGFP was kindly provided by Dr Meng Lin, School of Life Sciences, Sun Yat-Sen University. Cells were infected at various MOIs, as previously described ^26^.

### Real-time PCR

Total RNA was extracted using TRIzol reagent (Invitrogen) and subjected to reverse transcription using HiScript® III RT SuperMix (Vazyme). Real-time PCR was performed with a LightCycler® 480 SYBR Green I Master Mix kit (Roche). The primers used in the indicated gene array are listed in Table S1. The following primers were used for real-time PCR:

*TNFα*:

Forward 5’-CTCTTCTGCCTGCTGCACTTTG-3’

Reverse 5’-ATGGGCTACAGGCTTGTCACTC-3’

*IFNβ*:

Forward 5’-CCTACAAAGAAGCAGCAA-30-3’

Reverse 5’-TCCTCAGGGATGTCAAAG-30-3’

*IFIT1*:

Forward 5’-GCCTTGCTGAAGTGTGGAGGAA-3’

Reverse 5’-ATCCAGGCGATAGGCAGAGATC-3’

*IFIT2*:

Forward 5’-GGAGCAGATTCTGAGGCTTTGC-3’

Reverse 5’-GGATGAGGCTTCCAGACTCCAA-3’

*IL-6*:

Forward 5’-AGACAGCCACTCACCTCTTCAG-3’

Reverse 5’-TTCTGCCAGTGCCTCTTTGCTG-3’

*CCL20*:

Forward 5’-AAGTTGTCTGTGTGCGCAAATCC-3’

Reverse 5’-CCATTCCAGAAAAGCCACAGTTTT-3’

### Immunoprecipitation and immunoblot analysis

For immunoprecipitation, whole cell extracts were prepared after transfection or stimulation with appropriate ligands, followed by incubation for 1 h at 4°C with anti-GFP agarose beads (AlpaLife). Beads were washed 4 times with low-salt lysis buffer, and immunoprecipitants were eluted with 2x SDS loading buffer and then resolved by SDS-PAGE. Proteins were transferred to PVDF membranes (Millipore) and further incubated with the appropriate primary and secondary antibodies. The images were visualized using Odyssey Sa (LI-COR).

### Computer-based prediction and structural modeling

NSP8.pdb, MDA5-CARDs.pdb and K63-Ub.pdb were generated in SWISS-MODEL ^27^. MDA5-CARDs.pdb was input into ZDOCK-SERVER ^28^ as a receptor, while NSP8 or K63-Ub was input as a ligand for docking computation. MDA5-CARDs with NSP8.pdb and MDA5-CARDs with K63-Ub.pdb were the best fit prediction models chosen from the results. All the pdb files were processed and visualized with PyMOL (Schrödinger.

## Author contributions

X.L. and E.K. initiated the concept. Z.Y., X.L. and E.K. designed the experiments and analyzed the data. P.W. provided the reagents. Z.Y., X.Z., F.W. performed the experiments. Z.Y. and E.K. wrote the paper.

## Acknowledgments

We thank all the members of our laboratory for their critical assistance and helpful discussions. This work is supported by grants from the Natural Science Foundation of China (81671996 and 81871643) to E.K. and the Natural Science Foundation of China (81971928) to X.Li.

---

**Supplementary Figure.1.**
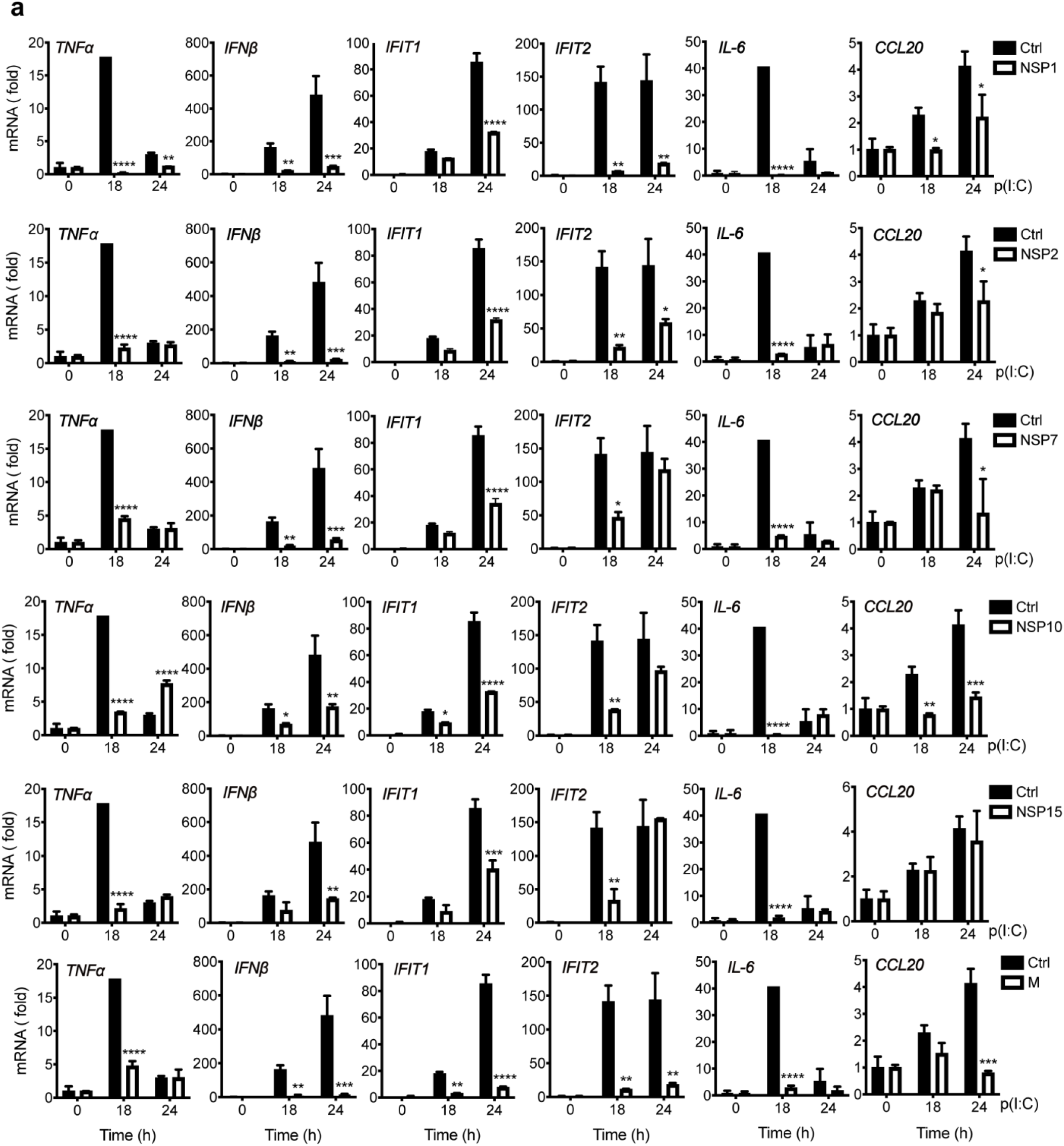
Screening results of SARS-CoV-2 nonstructural proteins in the regulation of type I IFN signaling. HEK293T cells were transfected with empty vector (Ctrl) or Flag-NSP plasmids. Twenty-four hours post transfection, cells were treated with poly(I:C) (5 μM) for the indicated time, and total RNA was subjected to RT-PCR analysis of TNF-α, IFN-β, IFIT1, IFIT2, IL-6 and CCL20 expression.

**Supplementary Figure.2.**
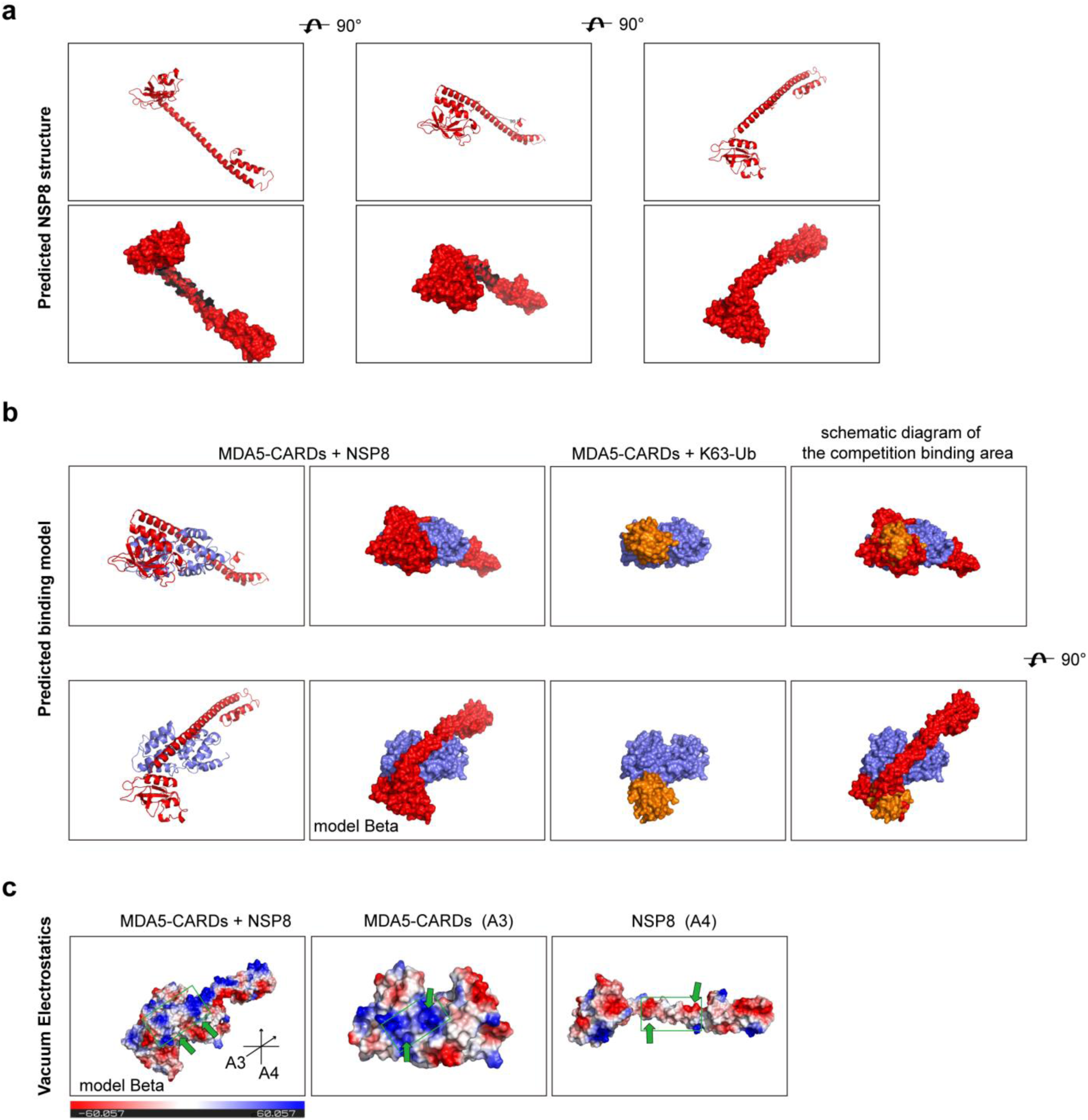
Predicted structure of NSP8 and its binding with MDA5 CARDs. a. The PDB structure of NSP8 was processed in PyMOL. The cartoon structure and surface structure are demonstrated. b. PDB structure of MDA5 CARD with NSP8 and MDA5 CARD with K63-Ub. PDB structures were input into the ZDOCK Server separately for docking calculations. The predicted binding models of MDA5 CARDs with NSP8 and MDA5 CARDs with K63-Ub were processed in PyMOL for demonstration. Model alpha simulates the protein surface. Red chain, NSP8; violet chain, MDA5 CARDs; brown chain, K63-Ub. c. Model alpha in (b) was subjected to vacuum electrostatics calculation in PyMOL. A3 and A4 indicate the viewing angle in the green frame. The green frame indicates the contact area demonstrated in A3 and A4. The scale bar indicates the range of vacuum electrostatics.

**Supplementary Table.1.**
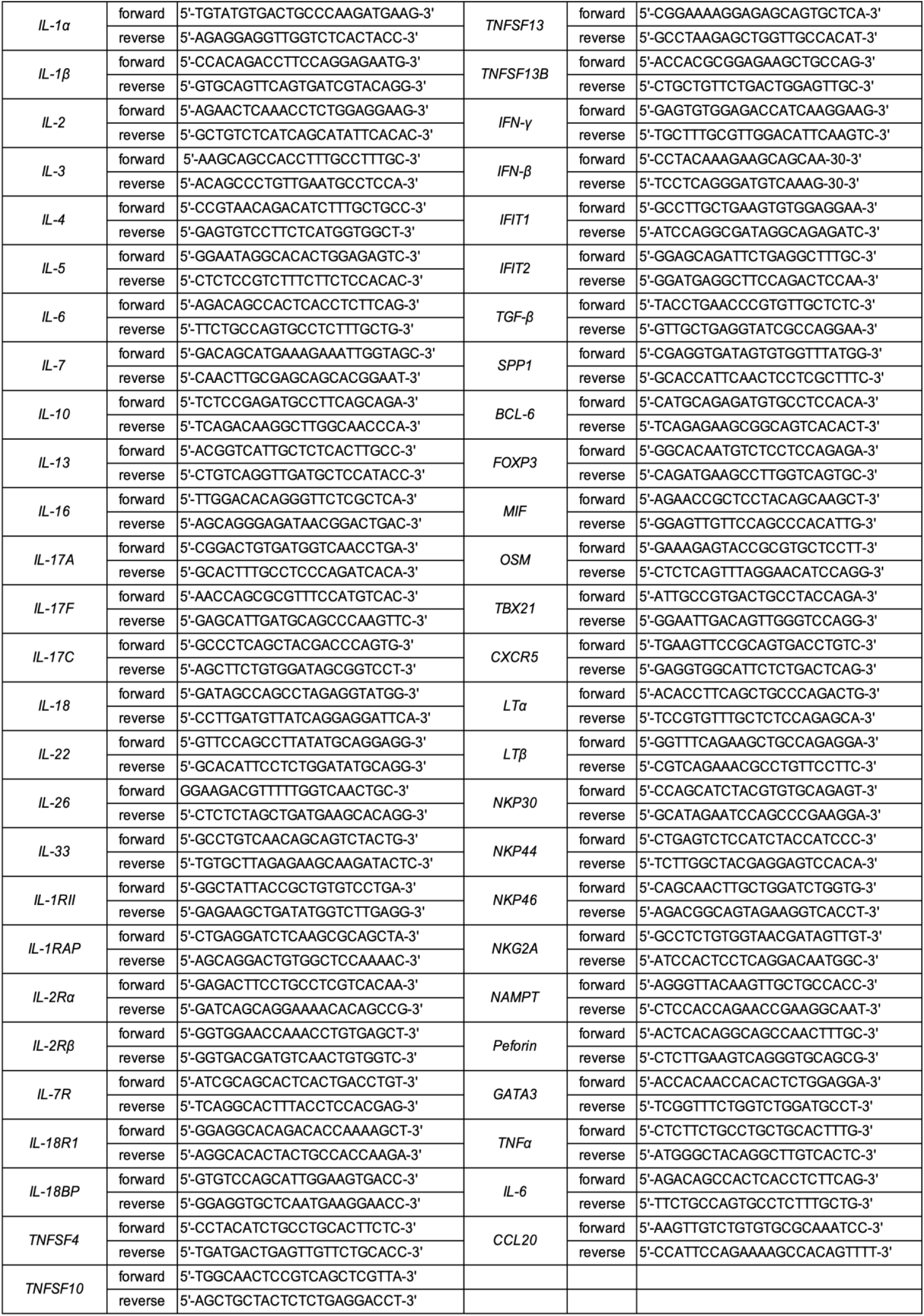
The primer pairs of the RT-PCR array. The sequences of primer pairs used in the RT-PCR array (Fig.5a) are listed.

